# Predicting Long and Short Duration Beta Bursts from Subthalamic Nucleus Local Field Potential Activity in Parkinson’s Disease

**DOI:** 10.1101/2023.09.22.558764

**Authors:** Bahman Abdi-Sargezeh, Sepehr Shirani, Abhinav Sharma, Philip A. Starr, Simon Little, Ashwini Oswal

## Abstract

Neural activities within the beta frequency range (13-30 Hz) are not stationary, but occur in transient packets known as beta bursts. Parkinson’s disease (PD) is characterized by the occurrence of beta bursts of increased duration and amplitude within the cortico-basal ganglia network. The pathophysiological importance of beta bursts is exemplified by the fact that they serve as a clinically useful feedback signal in beta amplitude triggered adaptive Deep Brain Stimulation (aDBS). Prolonged duration beta bursts are closely associated with motor impairments in PD, whilst bursts of shorter duration may have a physiological role. Consequently, we aimed to develop a deep learning-based pipeline capable of predicting long (*>* 150*ms*) and short (*<* 150*ms*) duration beta bursts from subthalamic nucleus local field potential (LFP) recordings. Our approach achieved promising accuracy values of 87% and 85.2% in two patients implanted with a DBS device that was capable of long-term wireless LFP sensing. Our findings highlight the feasibility of prolonged beta burst prediction and could inform the development of a new type of intelligent DBS approach with the capability of delivering stimulation only during the occurrence of prolonged bursts.

## 1 Introduction

Parkinson’s disease (PD) is a neurodegenerative condition charectrised by nigrostriatal dopamine deficit and the occurrence of stereotyped patterns of oscillatory synchrony within cortico-basal ganglia circuits [1], [2]. Disproportionate synchronization across the beta frequency band (13-30 Hz) characterizes the dopamine-depleted state in PD and is considered to be related to motor impairment [3]–[5]. Therapeutic approaches such as subthalamic nucleus (STN) deep brain stimulation (DBS) and dopaminergic medication help suppress basal ganglia beta oscillatory synchrony [2], [6].

Recent research suggests that beta activity related to PD emerges in brief periods called bursts [7], [8]. Although the mechanisms of beta burst generation are not fully understood, it is believed that bursts with long duration and high amplitude have a positive correlation with the degree of motor impairment in PD [9], leading to the beta activity being used as feedback in amplitude-responsive closed-loop DBS, where stimulation is delivered when beta amplitude rises above a specific threshold [6], [10], [11].

Beta bursts of short duration (less than 150 ms) are thought to play an important physiological role in sensorimotor processes [12]. In contrast, beta bursts of longer duration in PD are recognised to be pathological [7], [8]. This implies that neuromodulation strategies could potentially focus on selectively targeting these extended bursts while ignoring those shorter bursts that do not transition into a pathological state. These findings serve as a compelling incentive for the development of predictive algorithms aimed at identifying and precisely targeting pathological bursts.

Only one study has attempted to predict the occurrence of long-duration beta bursts [13]. This study did not however specifically address whether prolonged bursts could be differentiated from shorter bursts within the early phases (*<* 50*ms*) of burst onset. Addressing this question would be an important prerequisite for developing neuromodulation strategies that preemptively target long bursts.

Here we develop a deep neural network - based on Convolutional Neural Networks (CNNs) - to distinguish long- and short-duration beta bursts within 50ms of burst onset. This timely prediction presents an opportunity to deliver stimulation precisely when a long-duration burst commences. This approach holds the potential to offer several advantages over existing aDBS systems, including reduced battery consumption and fewer side effects.

## II Experiments and Methods

### A. Data Recording Details

Data from two patients with idiopathic PD were included in this study. Both patients underwent functional neurosurgery for the insertion of bilateral STN DBS electrodes. Recording STN LFP signals for a long term is significantly challenging and indeed a rare endeavor. Consequently, it is a common practice in these studies to analyze data from only two patients [13], [14].

STN LFP signals were recorded using the Summit RC+S bidirectional neural interface (Medtronic), which was attached to the quadripolar STN electrodes. This cutting-edge device boasts the capability to transmit neural data with an impressive sampling rate of up to 1,000 Hz to a Windows-based tablet located as far as 12 meters away, affording freedom of movement for users. Owing to this advantage, the data were recorded during normal activities of daily living in both awake and asleep states. Data were recorded in ON medication mode for three consecutive days while patients wore a Parkinson’s KinetiGraph (PKG) wrist watch [15]. These provided numerical scores for bradykinesia and dyskinesia every 2 min based on a 10-min moving average. The reader is referred to [16] for further information about data gathering.

Two bipolar channels located in each STN (L02, L13, R02, and R13 – L for the left and R for the right hemisphare) recorded LFP activities with a sampling rate of 250 Hz. The bipolar recordings were high-pass filtered at 1 Hz to remove DC offsets.

### B. Determination of Beta Peak Frequency

To determine the beta peak,we employed the short-time Fourier transform to derive the power spectrum of each bipolar contact. This spectral estimation was carried out using a Hamming window with a duration of 1 s and an overlap of 50%. The resulting spectra were then averaged across all time windows. Subsequently, these averaged spectra were visualized across a frequency range from 1 to 100 Hz, with a 1 Hz resolution (refer to Fig. 1a for an illustrative spectrum). For each participant, we selected a single bipolar LFP channel that exhibited the highest power peak within the beta frequency range (13-30 Hz). We refer to this channel as the “beta channel.” This channel was then chosen for further analysis.

**Fig. 1:**
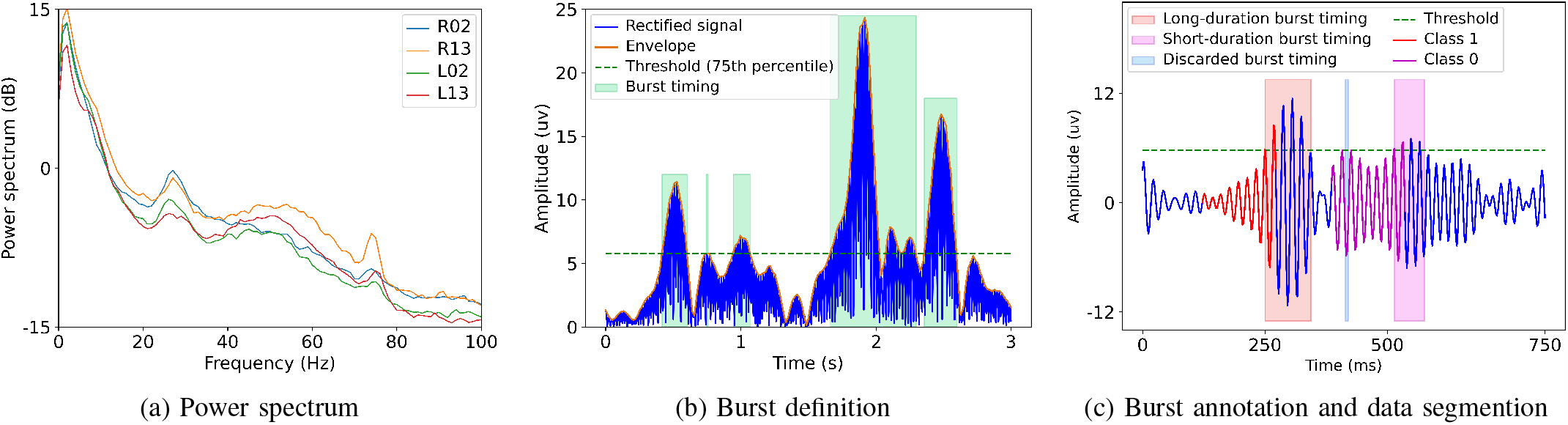
(a) demonstrates the power spectrum of right and left STN LFP signals of bipolar contact pairs 02 and 13. In this case (Patient 2), channel R02 has the greatest power peak within the beta range at a frequency of 27 Hz. R02 was therefore selected as the beta channel for Patient 2.(b) presents the rectified signal, its envolop, the threshold for burst definition, and burst timing. 75th percentile of beta envolop was used as the threshold. (c) shows the beta channel signal filtered ±3 Hz around beta peak frequency (for this case, from 24 to 30 Hz), long-duration burst timing (> 150*ms*), short-duration burst timing (> 50*ms* & > 50*ms*), discarded burst timing (< 50*ms*), and the data segments selected as Calss 0 and 1 respectively for short- and long-duration burst prediction.

### C. Definition of Long and Short Duration Bursts

To define the bursts, the beta channel of each patient was bandpass filtered within a ±3 Hz window centered on the beta peak frequency. To achieve this, we utilized a causal 6th order Butterworth filter. In this context, the use of a causal filter was crucial as it prevented any future signal components from affecting the current or previous filter outputs, which were utilized for prediction and analysis. Following bandpass filtering, the data were rectified before peak values were linearly interpolated to produce the beta amplitude envelope of the signal. Finally, we defined the onset and offset of each beta burst by identifying points where the beta amplitude envelope exceeded or fell below the 75th percentile of the amplitude envelope distribution as depicted in Fig. 1b.

Bursts lasting longer than 150 ms were categorized as “long bursts,” referred to as Class 1, while those with durations shorter than this threshold were classified as “short bursts,” referred to as Class 0. Bursts shorter than 50 ms were discarded from the analysis, depicted in Fig. 1c. This decision was based on the assumption that real-time data analysis typically involves the transmission of data in packets rather than on a sample-by-sample basis. Consequently, very short bursts were deemed unlikely to be reliably captured for prediction purposes. In addition, consecutive bursts with interburst intervals shorter than 50ms were merged together.

After defining burst timings, we proceeded to segment the time-domain LFP data, which would serve as the basis for training a neural network capable of reliably predicting whether a burst would be of long or short duration. Each segment was set to a length of 300 ms, spanning 250 ms before the onset of the burst and 50 ms after it occurred. In essence, we aimed to make predictions about the burst’s duration just 50 ms after its initiation. While this approach may not be ideal, it reflects the real-world conditions. When conducting real-time analysis, it’s essential to analyse the comming signal in a sliding window approach. The stride for updating the data within the window (the step size between consecutive windows) can be from 50 ms [13] to 400 ms [8]. Consequently, selecting a 50-ms stride is a logical and well-supported choice, as it aligns with established practices in the field.

### D. Split of Data to Trian, Validation, and Test Datasets

A total of 1307.7 and 1255.1 minutes of data were respectively gathered from patients 1 and 2, as outlined in TABLE I. The data were divided into training, validation (val) and test subsets for model evaluation. Multiple models were trained using the training dataset. Subsequently, the val dataset played a crucial role in determining the best-performing model among the trained ones. Once the best model was identified, it was evaluated using the unseen test dataset, providing a comprehensive assessment of its performance and generalization capabilities. Note that the traing and val/test datasets were selected from different recording days. This approach helps to ensure that the model doesn’t become overly specialized or biased towards specific recording conditions, making it more robust and adaptable to real-world scenarios where data can vary from day to day.

**TABLE 1:** Data details. The duration of recordings in minute and the number of longand short-duration bursts are illustrated.

| Patient | Duration (min.) |  |  |  | Num. of long bursts |  |  |  | Num. of short bursts |  |  |  |
| --- | --- | --- | --- | --- | --- | --- | --- | --- | --- | --- | --- | --- |
|  | Train | Val | Test | Total | Train | Val | Test | Total | Train | Val | Test | Total |
| 1 | 970.2 | 152.8 | 132.1 | 1255.1 | 36465 | 4648 | 4498 | 45611 | 36740 | 6248 | 4950 | 47938 |
| 2 | 1033.3 | 136.6 | 137.1 | 1307.0 | 40832 | 3553 | 3444 | 47829 | 34451 | 3741 | 3896 | 42088 |

The beta peak, beta channel, and the threshold for defining bursts for each patient were determined based the training dataset, which can provide a more comprehensive representation. The number of long- and short-duration bursts for each patient and dataset is illustrated in TABLE I.

### E. Prediction Network

The prediction network, shown in Fig. 2, was designed based on CNNs. First, the LFP signal was fed to the sequence of 1D convolution, rectified linear unit (ReLU) activation function, and max pooling. This series was repeated three times to capture deep temporal features of the signal. The output of last Max pooling was followed by a fully connected layer. At the end, a sigmoid function was employed to perform a binary classification, 0 (short-duration bursts) or 1 (longduration bursts).

**Fig. 2:**
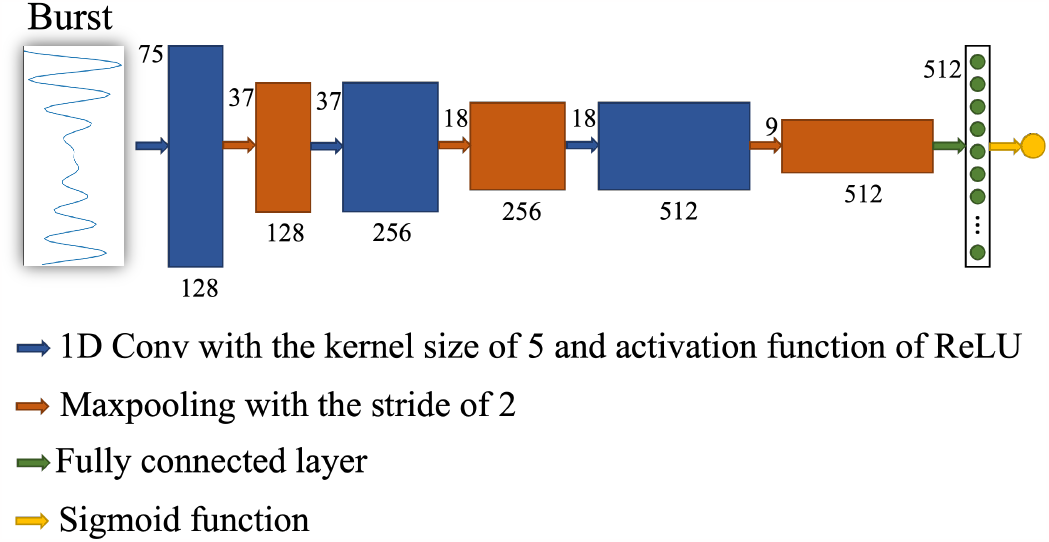
The proposed network architecture.

## III. Results

Accuracy, sensitivity (SEN), specificity (SPC), precision (PRC), the receiver operating characteristic curve (ROC), and area under the ROC curve (AUC) were obtained as the eveluation criteria, outlined in TABLE II and Fig. 3. ACC determines how well the model predicts long- and short-duration bursts and is defined as

**Fig. 3:**
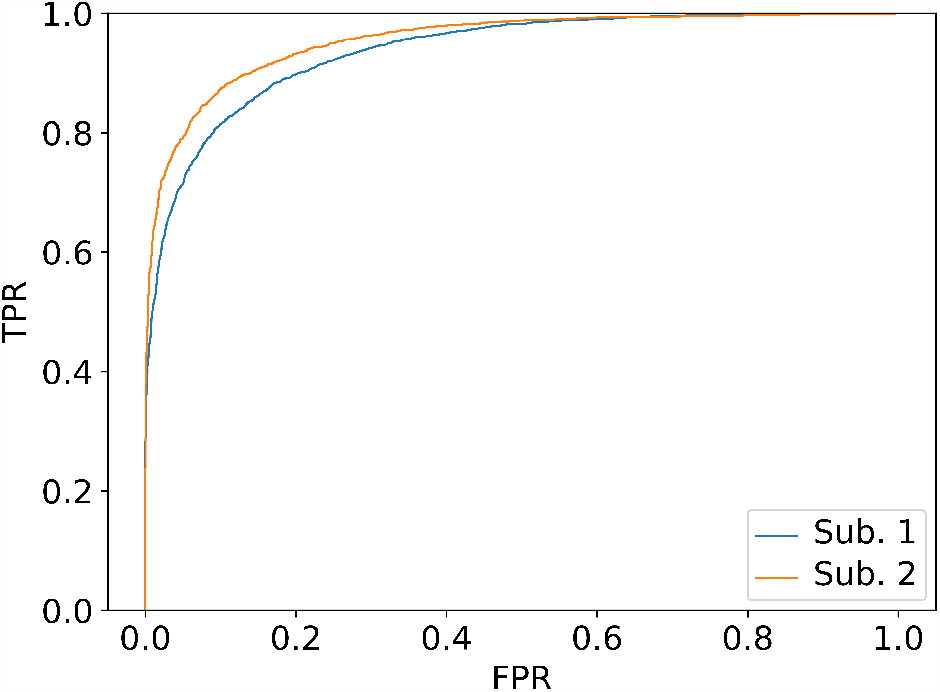
The ROC curve for patients 1 and 2.

**TABLE 2:** The performances of the network in distinguishing long and short bursts. ACC, SEN, SPC, and PRC are presented in percent (%).

| Patient | ACC | SEN | SPC | PRC | AUC |
| --- | --- | --- | --- | --- | --- |
| 1 | 85.7 | 85.2 | 86.1 | 84.8 | 0.93 |
| 2 | 88.6 | 86.5 | 90.5 | 89.9 | 0.95 |

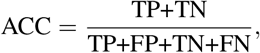

where TP is the abbrevation of true positive and shows the number of long-duration bursts predicted correctly by the pipeline, TN is the abbrevation of true negative and presents the number of short-duration bursts predicted correctly by the network, FP is the abbrevation of false positive and indicates the number of short-duration bursts that the pipeline incorrectly predicted as the long-duration bursts, FN is the abbrevation of false negative and illustrates the number of long-duration bursts that the pipeline incorrectly predicted as the short-duration bursts. SEN and SPC illustrate how well the model predicts respectively long- and short-duration bursts and are defined as

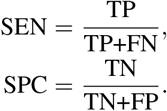

PRC shows the quality of the model in long-duration burst prediction and is defined as

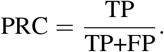

ROC and AUC are metrics that considers the trade-off between TP rate (TPR) and FP rate (FPR) at various threshold values.

The network achieved ACC values of 85.7% and 88.6% for subjects 1 and 2, respectively. In terms of SEN and SPC, the network provided respectively 85.2% and 86.1% for patient 1. On the other hand, for patient 2, it achieved SEN and SPC values of 86.5% and 90.5%, respectively. PRC values and AUC scores were higher than 84% and 0.93.

The ROC curve shown in Fig. 3 is a curve based on TPR against FPR. Achieving a high TPR comes at the cost of a high FPR. In real-world applications, we have the flexibility to adjust the threshold based on our specific problem requirements. By doing so, we can prioritize a high TPR, which is critical in our context as it minimizes the risk of missing long-duration burst predictions. This adjustment involves lowering the threshold of the sigmoid function to enhance SEN or TPR while accepting a trade-off of reduced SPC.

## IV. Conclusion

Here, we developed a pipiline based on deep learning to predict whether the duration of beta bursts will be long or short. The pipiline was applied to long-term STN LFP recordings of two PD patients. It achieved promising accuracy values of 85.7% and 88.6% respectively for patients 1 and 2. It also provided AUC scores of 0.93 and 0.95 for patients 1 and 2, respectively. These findings pave the way for the development of more intelligent and effective aDBS systems to improve the lives of PD patients.

